# Microbe-induced plant resistance against insect pests depends on timing of inoculation, but is consistent across climatic conditions

**DOI:** 10.1101/2023.09.07.556616

**Authors:** Oriana Sanchez-Mahecha, Sophia Klink, Michael Rothballer, Sarah Sturm, Wolfgang W. Weisser, Sharon Zytynska, Robin Heinen

**Affiliations:** Technical University of Munich, Terrestrial Ecology Research Group, Department for Life Science Systems, School of Life Sciences, Hans-Carl-von-Carlowitz-Platz, 85354 Freising, Germany; German Research Center for Environmental Health, Institute of Network Biology, Ingolstädter Landstraße 1, 85764, Neuherberg, Germany; University of Liverpool, Department of Evolution, Ecology, and Behaviour, Institute of Infection, Veterinary and Ecological Sciences, Crown Street, L69 7ZB, Liverpool, United Kingdom

**Keywords:** *Acidovorax radicis*, aphid suppression, bacteria-plant-insect interactions, barley plant, extreme climatic effect, herbivory, plant water availability, temperature extremes

## Abstract

1. To cope with abiotic and biotic stressors, plants have developed mutualistic associations with beneficial soil microbes, but little is known about how (extreme) abiotic conditions impact on microbe-induce resistance to insect herbivores.
2. Extreme temperatures are often accompanied by extremes in plant water availability, which together reduce plant growth and change plant physiology. There are potential consequences for increasing plant susceptibility to biotic stresses, and this poses a real challenge for plant productivity.
3. We evaluated how the effects of beneficial soil bacteria (*Acidovorax radicis*) on barley plant growth and resultant resistance against aphid infestation (*Sitobion avenae*) were impacted by a single heatwave event across a plant water availability gradient. We also tested if timing of bacterial inoculation (before or after the heatwave) affected bacteria-plant interactions on aphids.
4. We found that heatwaves affected plant biomass allocation from aboveground to belowground tissues. Inoculation with *A. radicis* led to reduction of aphid numbers, but depended on timing of inoculation, and led to stronger resistance when inoculations occurred closer to aphid infestation. Remarkably, microbe-induced resistance against aphids was consistent across heatwave and water availability treatments.
5. This study provides evidence that beneficial plant-bacteria interactions may represent a potential solution for sustainable agricultural practices to enhance plant growth and response to insect pests under climate change. Future field trials should investigate the consistency of beneficial effects reported here for a better understanding of multispecies interactions in the context of global change.

## Introduction

Plants have developed mutualistic associations with other organisms to mitigate stress. For instance, plants interact with rhizosphere microbes (Philippot et al., 2013; Vandenkoornhuyse et al., 2015), including beneficial bacteria. These bacteria can increase productivity, and enhance tolerance to abiotic factors (Bakker et al., 2018; Dimkpa et al., 2009; Van Oosten et., 2017) and often invoke microbe-induced resistance to pathogens and insects in plants (Heinen et al. 2018; Pineda et al., 2012; Pineda et al., 2017; Rashid & Chung, 2017; Sanchez-Mahecha et al., 2022). For example, soil bacteria of the genus *Bacillus* have been shown to reduce aphid populations on *Arabidopsis* (Rashid et al., 2017) and broccoli plants (Gadhave et al., 2016a) by inducing plant defence responses against the herbivores, or by attracting natural enemies to the host plant. Plant-bacterial interactions are gaining attention as a potential solution to increase stress tolerance in crops (Backer et al., 2018; Bakker et 2020; Timmusk et al., 2014). Despite this, knowledge of beneficial plant-microbe interactions under variable environmental conditions - especially for understanding high-order ecological interactions - remains rudimentary (Heinen et al., 2018; Zytynska, 2021).

Over the past decades, extremes and variability in temperature and precipitation have been common and are expected to occur more frequently under predicted climate change scenarios (IPCC, 2022). Heatwaves are defined as prolonged intervals (typically > 3 days) of elevated temperatures above a certain threshold (typically > 30°C), although local definitions may differ between countries (IPCC, 2022; Perkins & Alexander, 2013). Heatwaves often occur in combination with extreme precipitation regimes, such as drought or flooding, and are simultaneously experienced by plants and other organisms. However, these abiotic factors have mainly been studied independently from each other, and only a few studies have considered extreme temperature effects on plants in combination with other abiotic factors (De Boeck et al., 2016; Duan et al., 2016; Lamaoui et al., 2018; Marchin et al., 2021). Such combined conditions can cause large-scale ecological and economic problems, including catastrophic crop failure (Beillouin et al., 2020), soil degradation, local pest outbreaks, and may influence ecological interactions (Harvey et al., 2020; Meisner et al., 2013; Xu et al., 2016). Studying ecological interactions under multiple interacting climate stressors is direly needed to increase our understanding of interactions in a changing world (Rillig et al., 2019).

Climatic extremes can strongly impact plant-mediated interactions by influencing any of the involved organisms directly, or indirectly, by producing changes in the plant host that can have consequences for associated interactions. Heatwaves alone or in a combination of extremes in water availability, such as low water availability, negatively affect plant growth and photosynthetic activity (Zandalinas et al., 2018) by increasing the plant investment in transpiration processes to cool down vital organs during high-temperature periods (Lamaoui et al., 2018). This generally leads to reduced primary plant productivity (Christmann et al., 2007; Takahashi et al., 2020; Tombesi et al., 2015) with consequences for plant nutritional quality and investment in defence-related pathways (Fraser & Chaple, 2011; Zandalinas et al.,2017). Under heat stress, for example, plants show specific molecular and physiological responses such as the production of secondary metabolites (Zandalinas et al., 2017), which have been associated with plant survival and acclimation to environmental stress conditions and have also been associated with increased plant defences (Fraser & Chaple, 2011; Rashid et al., 2018). Plant physiological changes during heatwaves or extremes in water availability might influence its response to other biotic stress factors, such as herbivores, by increasing plant vulnerability to feeding insects (Harvey et al., 2020; Showler et al., 2013). Low water availability, for example, reduces plant cell water content and concentrates shoot nitrogen levels, which has been shown to result in positive bottom-up effects on herbivores and higher vulnerability for the plant (Rivelli et al., 2013; Showler et al., 2013), but the effect depends on the duration and severity of the water stress (Rivelli et al., 2013). Moreover, increased temperature and extremes in water availability during heatwave periods can also impact the soil microbiome (Meisner et al., 2018; van der Voort et al., 2016), potentially causing physiological changes in the plants that grow in these heat-exposed soils (Rubin et al., 2018) and as a cascading effect disturb plant-microbe and higher ecological interactions (Abarca & Lill, 2015; Schwartzberg et al., 2014; Ward et al., 2019). However, little is understood about the potential effects that combined climatic stressors may leave on plants and their associations with soil microbes, and how these associations affect plant ecological interactions in the future.

Here, we performed a climate chamber experiment to test the effects of a heatwave event across a gradient of plant water availability on interactions between beneficial bacteria (*Acidovorax radicis*), barley plants (*Hordeum vulgare*), and cereal aphids (*Sitobion avenae*). The aims of the present study were (i) to understand how interacting climatic factors (i.e., heatwaves and water availability) impact on plant performance and resistance against aphids, (ii) to understand how these climatic conditions may affect interactions between beneficial soil bacteria and plants, with consequences for microbe-induced resistance against aphids, and (iii) to understand how timing of inoculation (i.e., before or after the heatwave event) would affect microbe-induced resistance. Specifically, we tested the following hypotheses. (1) Heatwaves will reduce plant growth and chlorophyll content (a proxy of plant health), exacerbated under low water availability treatment. (2) Heatwaves will negatively affect plant resistance against aphids, allowing increased aphid growth rates, exacerbated under low water availability. (3) Pre-heatwave bacterial inoculation will help plants cope with a post-heatwave aphid infestation, but efficacy will be negatively affected by the heatwave, especially under low plant water availability. (4) A post-heatwave bacterial inoculation will increase microbe-induced resistance to aphids, and reverse potential negative effects caused by the heatwave.

## Materials and methods

### Experimental design

A full-factorial experiment was conducted in the Technical University of Munich Model Ecosystem Analyser (TUM*mesa*) climate chamber facilities in Freising, Bavaria, Germany. For this study, barley plants (*Hordeum vulgare*) of the cultivar Scarlett (Saatzucht Breun GmbH, Herzogenaurach, Germany) were selected. Barley is grown in many temperate areas of the world, and is economically highly relevant. Similar to many other cereal crops, barley plants experience strong colonization by aphids over the summer season, and environmentally sustainable solutions are highly sought after. Previous work revealed that the selected cultivar is sensitive to rapid aphid colonization, and that *Acidovorax radicis* bacteria provided consistent microbe-induced resistance against aphids (Sanchez-Mahecha et al., 2022).

Plants were exposed to two abiotic treatments: (1) a heatwave treatment (two levels: ambient/heatwave), and (2) a plant water availability gradient (four levels: minimally-watered 17.5%, moderately-watered 25%, optimally-watered 32.5%, over-watered 40% soil volumetric water content, respectively) and one additional biotic treatments: (3) an *Acidovorax radicis* N35e rhizobacterial inoculation (four levels: control (without bacteria), pre-heatwave inoculation, post-heatwave inoculation and both pre-and post-heatwave (double) inoculation). This resulted in 32 treatment combinations that were replicated 10 times (320 plants), from the 320 plant 3 were discarded because of poor growing features (<15 cm or almost dried < 20 chlorophyll content SPAD units). All plants were exposed to aphids (*Sitobion avenae*).

The experiment was executed over two runs (temporal blocks). Four climate chambers were used in each run to simulate the heatwave (two chambers with a heatwave and two with ambient conditions per run). Within the heatwave treatments, soil moisture and rhizobacterial treatments were performed at the individual pot level, and the 10 replicates of each treatment combination were divided over the two runs (Fig. 1). The chamber treatment assignment was switched between chambers between runs, to correct for potential chamber effects. Pots were placed in individual trays for watering and were completely randomised inside the chamber every week.

**Figure 1:**
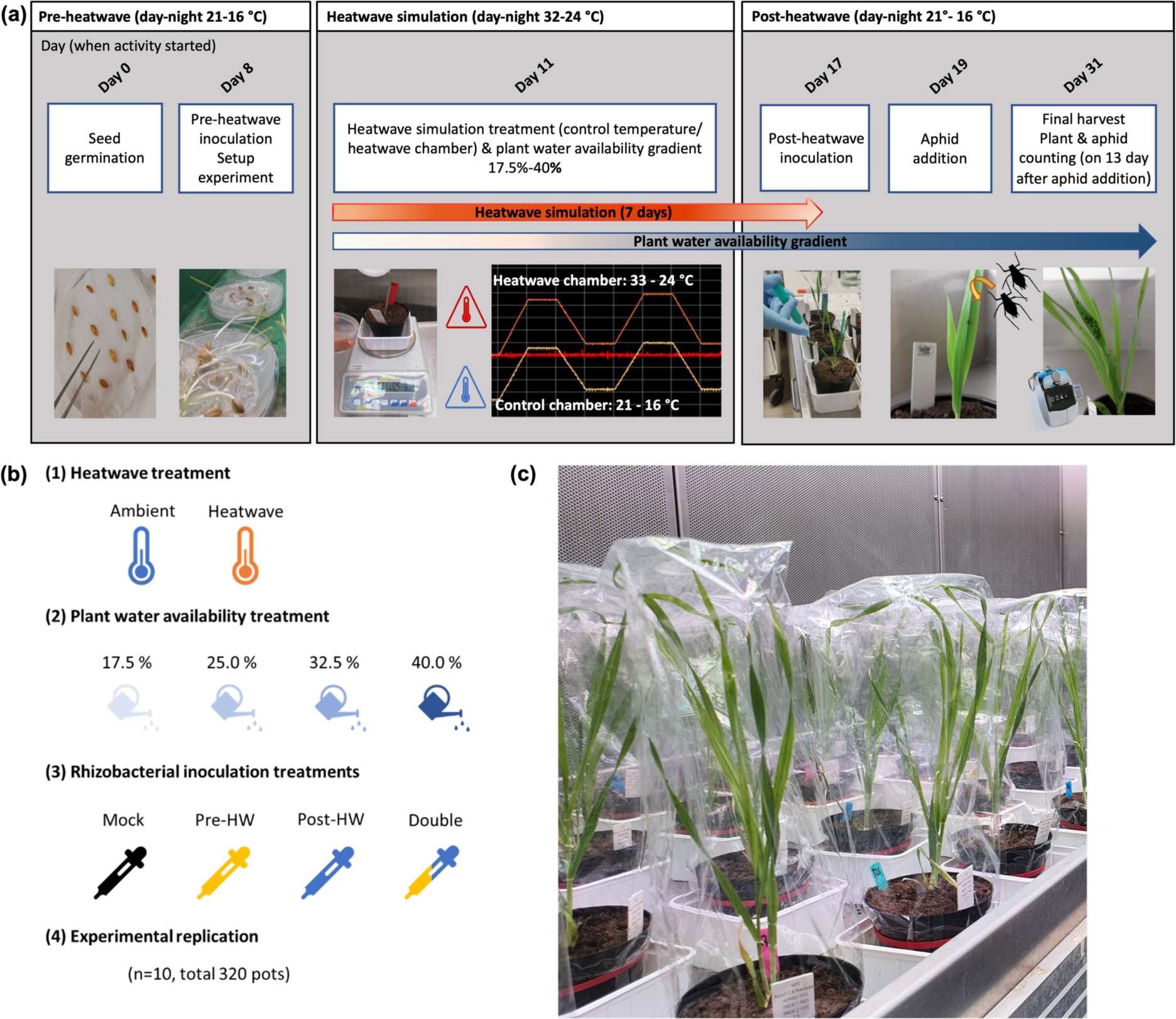
a) Overview of the experimental timeline, divided into the different experimental stages (i.e. pre-heatwave, heatwave, post-heatwave) adopted in the ecological experiment. Timing of experimental procedures is always indicated in days post-germination. The arrows in the heatwave and plant availability gradient show the time period of the treatment in days. b) Overview of the experimental design including levels of each experimental factor, i.e., heatwave, plant water availability, and rhizobacterial inoculation treatments. c) Picture of the experimental plants after aphid infestation.

### Heatwave treatment

The heatwave treatment was simulated based on local definitions for Germany, although similarity exists among heatwave definitions. As Germany is an important producer of barley crop, and heatwaves occur regularly, local definitions are relevant guidance for this model system. We define heatwaves as a period of high temperatures that exceed the maximum day-time threshold temperature of 30 °C for at least three consecutive days (Tomczyk & Sulikowska, 2018). Our treatment had two levels, heatwave and ambient (Fig. 1).

For the two growth chambers that were assigned to the ambient conditions, the temperature was kept at daily cycles of 21/16 °C day/night temperatures, light intensity ∼500 µmol/m²s,16:8 L:D photoperiod and RH 60% during the whole experiment. In the other two chambers that had a heatwave simulation, the temperature started with the same temperature settings as the ambient chambers, 21/16 °C. However, eleven days post-germination, the temperature was gradually increased for two days until it reached 33/24 °C day/night temperatures; this heatwave temperature was then kept for three consecutive days (Fig. 1). Subsequently, the temperature was reduced again gradually for two days until it reached the ambient temperature (21 °C/16 °C) that remained until the termination of the experiment.

### Plant water availability gradient treatment

We created a gradient of plant water availability with four levels: 17.5%, 25%, 32.5% and 40% volumetric water content (vwc), respectively. Target watering weights were determined for each of the four levels of the gradient. First, the soil dry mass of a filled pot including the watering tray was determined. To this, we added the weight of the respective target water volume (%) for each gradient. The plant water availability gradient was maintained daily by watering each plant to the desired weight on a balance. In the experiment, the levels of the plant water availability gradient were considered as follows: minimally-watered plants (17.5% vwc), moderately-watered plants, (25% vwc), optimally-watered plants (32.5% vwc) and excessively-watered watered plants (40% vwc). In minimally-watered plants the soil was always dried-out on the surface, on the moderately- and optimally-watered plants the soil was dry to mildly humid, and on excessively-watered plants, the soil was wet before the daily watering.

### Bacterial inoculum

In this study we used the beneficial rhizobacterium *Acidovorax radicis* that is known to induce suppressive effects against pest aphids, as well as protective effects against various abiotic stressors (Zytynska et al., 2020; Sanchez-Mahecha et al., 2022). *Acidovorax radicis* colonises plant roots by forming biofilm-like structures on the root surface (Han et al., 2016). Biofilm production has been reported to increase plant water retention in other bacterial species, which might also aid in the plants’ physiological recovery under stress conditions (Timmusk & Nevo, 2011; Valliere et al., 2020). The *Acidovorax radicis* and control bacteria solutions (without bacteria) were prepared and inoculated twice per run before and after the heatwave event. The bacteria were grown on NB agar plates, later the bacterial lawn was collected and re-suspended in 10 mM MgCl_2_ with Tween20 (100 µl/L) to an optical density adjusted to OD600 = 1.5 in the first preparation [approx. 10^8 colony-forming units (cfu/ml)] and OD600 = 0.5 in the second preparation (approx. 10^7 cfu/ml). In the second bacteria inoculation preparation, the concentration was reduced to OD600 = 0.5, since in the first inoculation the plant seedling roots were exposed to the bacteria solution for 1 hour only, while in the second inoculation, the bacteria were added permanently directly to the soil close to the plant roots. Therefore, to avoid bacterial overpopulation and create similar conditions in plant exposure to bacteria, the concentration was reduced in the second inoculation. The controls were inoculated with a solution that only contained 10 mM MgCl2 + Tween20 (100 µl/L).

### Insects

Cereal aphids (*Sitobion avenae* L; genotype ‘Fescue’) were reared on barley cultivar Chanson (Ackermann Saatzucht GmbH) in a growth chamber (Temperature 21 ± 1 °C, relative humidity (RH) 60%, and 16:8 L:D photoperiod). Aphids were obtained from clonal colonies that have been kept in low densities on barley cultivar Chanson for several years at the Terrestrial Ecology Research Group, Technical University of Munich. Cereal aphids (Hemiptera: Aphididae) are relevant economic pests because they produce significant plant yield losses annually by reducing the plant growth (Tatchell 1989; Larsson 2005; Dedryver, Le Ralec & Fabre 2010), and by transmitting important plant viruses as vectors (Kamphuis et al., 2013). Aphids, being parthenogenetic and viviparous organisms can be highly prolific and show exponential growth (Powell et al., 2007). Additionally, under present climate change scenarios, and need to phase out pesticides, it will be increasingly challenging to predict and manage insect pest populations, and therefore sustainable management solutions are needed.

### Experimental procedure

Before germination, barley seeds were surface-sterilised with 5% hypochlorite solution and thoroughly rinsed with water. After that, seeds were germinated between filter papers for eight days in a dark cabinet at room temperature. Five days post-germination, pots were filled with 90 g of potting soil (Floradur multiplication substrate, Floragard; 140N, 80P, 190K, pH 6.1), and each pot was watered with 20 ml directly on the soil to facilitate the bacterial treatment by preventing soil drying and creating similar soil moist conditions between pots. Eight days post-germination, barley seedlings with approximately uniform root and shoot sizes were divided into two groups for the bacterial inoculation: (1) pre-heatwave inoculation with *A. radicis*, and (2) control plants exposed to control solution. Seedling roots were soaked in these solutions for 1 hour. One seedling was planted per pot, shoots were measured, and plants were allocated to the corresponding climate chamber. Plants were watered directly on the soil with 30 ml of tap water.

Eleven days post-germination, the water availability gradient treatment started and lasted until the end of the experiment, including the heatwave treatment that lasted for seven consecutive days (Fig. 1). After the heatwave treatment, on day 17, the post-heatwave bacterial inoculation took place. Plants were *a priori* assigned to two groups: post-heatwave bacterial inoculation or post-heatwave control inoculation.

Two days after the second bacterial inoculation, two sub-adult aphids (4th-instar) were added to the base of all plant shoots. After this step, all plants were covered with an air-permeable cellophane bag (HJ Kopp GmbH, Germany) and secured with an elastic band to prevent aphid escape. Total aphid numbers were counted on day 13 after aphid addition, corresponding to day 31 post-germination in the experiment, and on this day the final destructive harvest was conducted. The following variables were measured: shoot length, number of leaves and stems, shoot fresh weight, chlorophyll content (with Konica Minolta SPAD-502), shoot dry biomass (after drying the samples for four days at 45 °C), root biomass (after washing and drying for eight days at 45 °C).

### Statistical analysis

All the analyses were performed in RStudio version 1.4.1106 (RStudio Team, 2021) using R version 4.0.5 (R Core Team, 2021). Plant and aphid data were analysed with linear models. The plant response variables were shoot biomass, root biomass, and chlorophyll content (day 31 post-plant germination). The insect response variable was aphid colony size, specifically the number of aphids on day 13 after aphid addition (corresponding to day 31 post-plant germination). Plant and insect full models contained the main effects (‘heatwave’, ‘plant water availability gradient’, rhizobacteria inoculation), and their interactions. As blocking factors, we included ‘run’, and ‘climate chamber’ in the model to control the variation between the temporal separation of the two runs and potential climate chamber effects. After running the model, a simplification step was implemented using a backwards stepwise method using the step() function in R and by excluding one by one the least significant interaction term until getting the minimal adequate model result. Model output can be found in Table 1, and AIC values can be found in Table S2. All data and model residuals were checked for normality and heteroscedasticity through diagnostic plots, and data were transformed as necessary (i.e., log(root biomass + 1); log(shoot.root_ratio+1)).

**Table 1.**
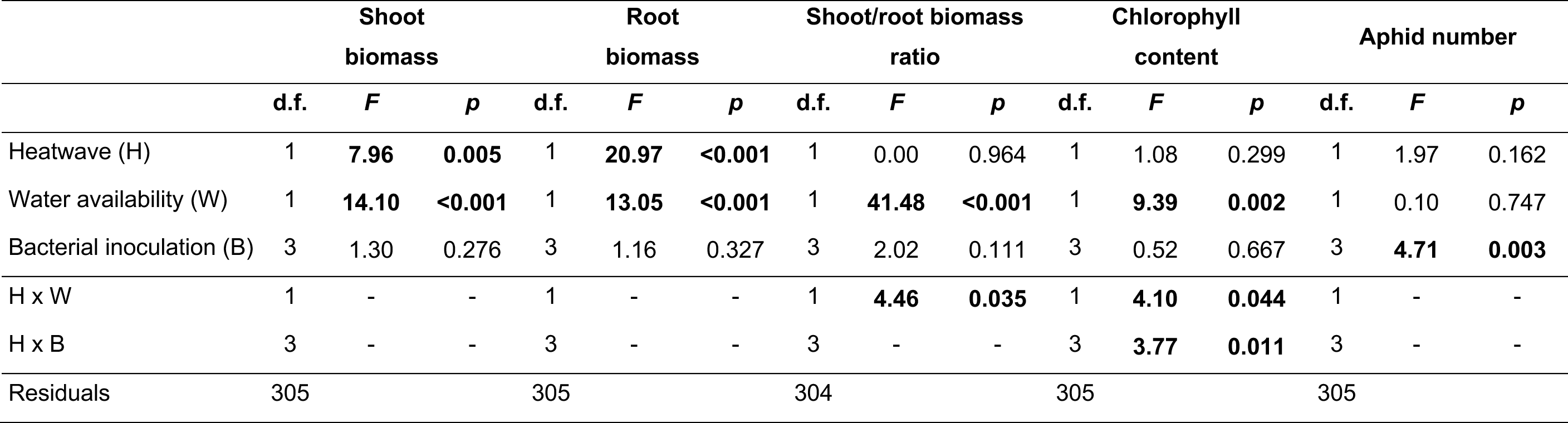
Summary of standard linear models for the plant and aphid response variables. Abbreviations on the table are as follows: Heatwave (H), Water availability (W) and Bacterial inoculation (B). *Note:* Data were transformed as necessary to approach model assumptions ([log (root biomass + 1)]; [log(shoot. root_ratio+1)]). “Bold values” show significant p values (<0.05), from minimal adequate models. Dashes indicate terms removed after model simplification.

|  | Shoot biomass |  |  | Root biomass |  |  | Shoot/root biomass ratio |  |  | Chlorophyll content |  |  | Aphid number |  |  |
| --- | --- | --- | --- | --- | --- | --- | --- | --- | --- | --- | --- | --- | --- | --- | --- |
|  | d.f. | *F* | *p* | d.f. | *F* | *p* | d.f. | *F* | *p* | d.f. | *F* | *p* | d.f. | *F* | *p* |
| Heatwave (H) | 1 | **7.96** | **0.005** | 1 | **20.97** | **<0.001** | 1 | 0.00 | 0.964 | 1 | 1.08 | 0.299 | 1 | 1.97 | 0.162 |
| Water availability (W) | 1 | **14.10** | **<0.001** | 1 | **13.05** | **<0.001** | 1 | **41.48** | **<0.001** | 1 | **9.39** | **0.002** | 1 | 0.10 | 0.747 |
| Bacterial inoculation (B) | 3 | 1.30 | 0.276 | 3 | 1.16 | 0.327 | 3 | 2.02 | 0.111 | 3 | 0.52 | 0.667 | 3 | **4.71** | **0.003** |
| H x W | 1 | - | - | 1 | - | - | 1 | **4.46** | **0.035** | 1 | **4.10** | **0.044** | 1 | - | - |
| H x B | 3 | - | - | 3 | - | - | 3 | - | - | 3 | **3.77** | **0.011** | 3 | - | - |
| Residuals | 305 |  |  | 305 |  |  | 304 |  |  | 305 |  |  | 305 |  |  |

## Results

### Heatwave and water availability effects on plants

Heatwave conditions and plant water availability had significant main effects on most plant variables (Table 1). Heatwave conditions reduced shoot biomass compared to ambient conditions, and shoot biomass generally increased with increasing plant water availability (Fig. 2a, Table 1). An opposite tendency was observed for root biomass, where heatwave conditions increased root biomass compared to ambient conditions while higher plant water availability reduced it (Fig. 2b, Table 1). The shoot-root ratio increased with more water availability, but this effect was observed mostly on plants that were not exposed to heatwave conditions. This suggests that an increase in water availability increased the biomass allocation from the root to the shoot but only on ambient plants, while on heatwave-exposed plants, the observed change in the shoot-root ratio was smaller (Fig 2c, Table 1). Chlorophyll levels were affected by plant water availability, where chlorophyll decreased with increasing water availability (Table 1). However, the effect of water availability on chlorophyll interacted with the heatwave treatment, as the decrease was observed under ambient conditions, but less so under heatwave conditions (Fig. 2d, Table 1).

**Figure 2:**
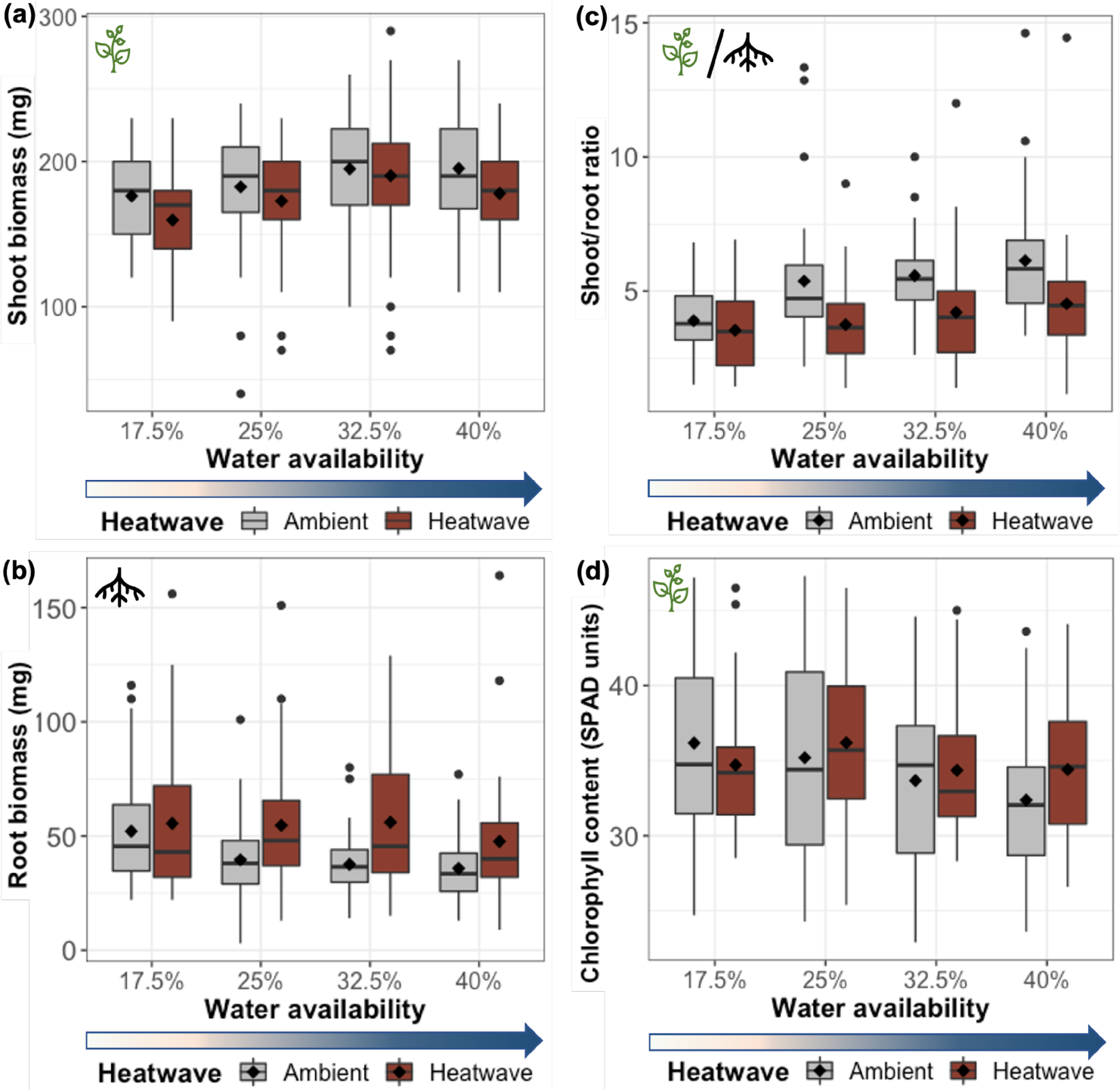
Heatwave (H) and plant water availability (W) effects on barley plant (*Hordeum vulgare*). The panels show effects on (a) shoot biomass (H: F_1,305_ =8.0, *P* = 0.005; W: F_1,305_ = 14.1, *P* < 0.001), (b) root biomass (H: F_1,305_ = 21.0, *P* < 0.001; W: F_1,305_ = 13.0, *P* < 0.001), (c) shoot-to-root ratio (W: F_1,304_ = 41.5, *P* < 0.001, H x W: F_1,304_= 4.5, *P* = 0.035), (d) late chlorophyll content (measured on day 31 post-germination, W: F_1,305_ =9.4, *P =* 0.002; H x W: F_1,305_= 4.1, *P* = 0.044). Boxes represent median values with upper and lower quartiles, and whiskers represent 1.5 x the interquartile range. The diamond symbol in the boxes indicates the group mean. Individual points represent outlier data points (n=10 replicates).

### Heatwave and water availability effects on insects

Heatwave and water availability, or their interactions, did not affect aphid colony size, suggesting that plant exposure to heatwave conditions prior to aphid infestation does not leave long-term effects on plants that affect aphids arriving later on the plant (Table 1).

### Rhizobacteria (*Acidovorax radicis*) inoculation timing effects on aphid colony growth

Inoculation with *Acidovorax radicis* resulted in microbe-induced plant resistance that significantly reduced aphid numbers, independently of the water plant availability or the heatwave conditions (Fig. 3, Fig. S1; Table 1). Interestingly, the effect of *A. radicis* inoculation differed between the four inoculation types, and posthoc Tukey tests revealed that significant suppressive effects compared to no inoculation controls were only observed on the plants that had an after the heatwave inoculation treatment (i.e., post-HW and double inoculation treatments (Fig.3, Table 1). A separate model treating pre-heatwave and post-heatwave inoculations as separate factors showed that both inoculations resulted in aphid suppression, but that the effect of post-heatwave inoculations was stronger (pre: *P* = 0.029, post: *P* = 0.004, resp., Table S1).

**Figure 3:**
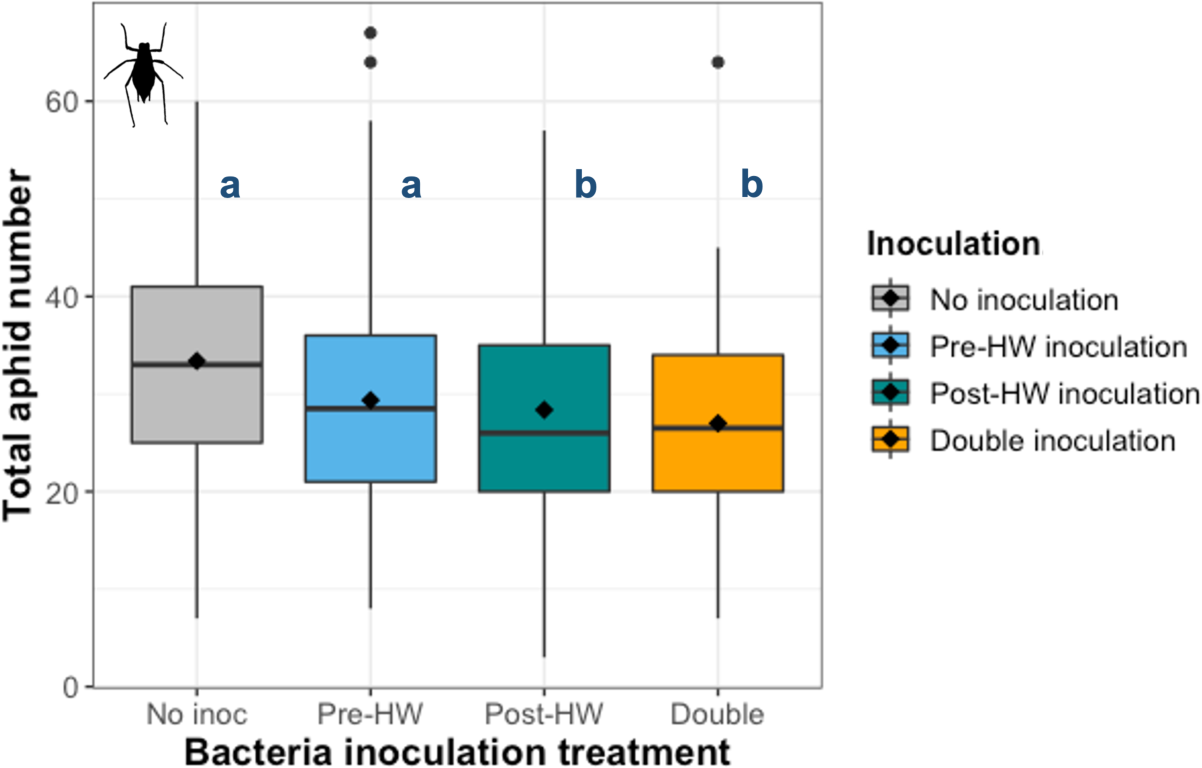
*Acidovorax radicis* bacterial inoculation effect on cereal aphid (*Sitobion avenae*) growth (F_3,305_ = 4.7, *P* = 0.003). Boxes represent median values with upper and lower quartiles, and whiskers represent 1.5 x the interquartile range. The diamond symbol in the boxes indicates the group mean. Individual points represent outlier data points (n=10 replicates), and the letters above the boxes represent significant differences (*P*<0.05) between groups estimated from marginal means EMMs, Tukey method, calculated with R version 4.0.5, package emmeans (pre-heatwave inoculation *P* = 0.104, post-heatwave inoculation *P* = 0.032, double inoculation *P* =0.002).

### Interactive rhizobacterial inoculation and climatic effects on plant performance

*Acidovorax radicis* bacterial inoculation did not alter plant biomass, plant shoot-to-root ratio, or chlorophyll content, and we observed no interactive effects with climatic factors in terms of plant growth. However, we observed an interactive effect between bacterial inoculation and heatwave treatment on chlorophyll content. Generally, chlorophyll content was slightly higher in the bacterial inoculation treatments under heatwave conditions than under ambient conditions, except in the post-heatwave inoculation treatment where the chlorophyll content was higher under ambient conditions than under heatwave conditions (Table 1; Fig. 4).

**Figure 4:**
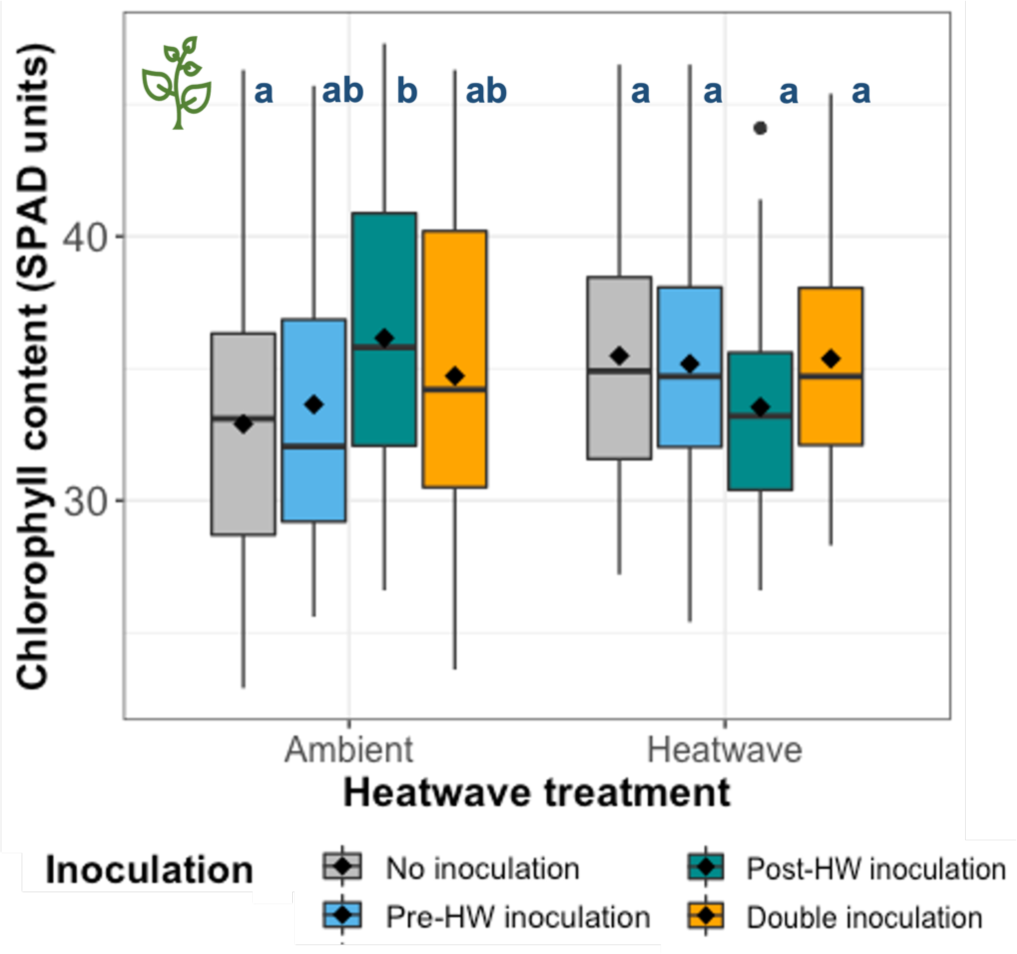
Effect of bacterial inoculation *(Acidovorax radicis)* on barley plant (*Hordeum vulgare*) chlorophyll content (SPAD units) across heatwave (H) treatments (H x B: F_3,305_ = 3.8, *P*=0.011). Boxes represent median values with upper and lower quartiles, and whiskers represent 1.5 x the interquartile range. The diamond symbol in the boxes indicates the group mean. Individual points represent outlier data points (n=10 replicates), and the letters above the boxes represent significant differences (*P*<0.05) between groups estimated from marginal means EMMs, Tukey method, calculated with R version 4.0.5, package emmeans. (Ambient conditions: No inoculation vs post-heatwave inoculation *P* = 0.023).

## Discussion

We evaluated the effect of heatwave conditions on the effects of a beneficial microbe on plant growth and microbe-induced plant resistance to aphids, across a plant water availability gradient. Heatwave conditions and water availability acted additively to shift the growth allocation from above to belowground tissues resulting in a negative effect on the shoot-to-root ratio with decreasing water availability. Bacterial inoculations decreased aphid numbers on the plants, with the strongest significant aphid suppression occurring in the post-heatwave bacterial inoculations. This suggests that an inoculation closer to the aphid infestation could provide stronger beneficial effects on the plant, or could provide reinforcement of microbe-induced resistance. Moreover, heatwaves and water availability did not interact with the microbial inoculation treatment to alter aphid suppression effects. Our findings underpin the role of beneficial microbes in providing consistent plant protection under abiotic or biotic stresses (Frew et al., 2022; Pieterse et al., 2016; Pineda et al., 2017).

### Heatwaves and plant water availability shift barley biomass allocation

Plant biomass allocation was strongly affected by heatwave exposure. Under heatwave conditions plants allocated biomass from aboveground to belowground parts, which aligned with our expectations. A likely explanation is that under heat exposure, plants reduce evapotranspiration and primary plant productivity via a reduction in shoot biomass (Flexas et al., 2004; Zandalinas et al., 2016). In addition, plants might invest their resources in their root system to maximize water absorption enabling survival in hot and dry conditions (Ober & Sharp, 2007; Sicher et al., 2012). Our results align with previous studies that also found that drought conditions negatively affected aboveground plant tissues and had a positive effect on root growth (De Bobadilla et al., 2017; Lamaoui et al., 2018; Shrap et al., 2004; Singh and Reddy, 2011; Zhao et al., 2017). Our results demonstrate the clear combined effect of heatwaves and plant water availability on above- and belowground biomass.

### Heatwaves and plant water availability did not affect aphid colony growth

Contrary to our hypotheses, climatic factors did not affect subsequent aphid colonies on plants, indicating that plant resistance against aphids was not altered by the climatic environment that the plants were exposed to. Various studies have shown physiological changes in plants under heatwave stress (Zandalinas et al., 2018; De Boeck et al., 2016; Duan et al., 2016; Lamaoui et al., 2018), and we did detect effects on biomass allocation; however, since our plants were infested with aphids after the heatwave treatment we conclude that any plant biochemical changes due to climate effects are short-lived. It is interesting that plant water availability, which was continuously maintained throughout the study, did not affect aphid colony growth. Given that aphids on the one hand depend on water and turgor for feeding (Huberty & Denno, 2004), but on the other hand can also be affected by concentration effects caused by water stress (Beetge & Krüger, 2019), it is surprising that aphids performed equally well on all levels of water availability. Although it could be argued that the lowest level (17.5% vwc) of water availability does not equate drought, we emphasize that plants used up water between waterings, with levels dropping below 10% vwc, and plants showed clear signs of water stress in terms of growth (Fig. 2) and low turgor pressure. As our experiment was executed in the early stages of plant growth and aphid colonization, it is more plausible that effects would manifest at later stages, but this would require further experimental efforts to test.

### *Acidovorax radicis* invokes microbe-induced resistance, which was influenced by the timing of inoculation but not by climatic factors

Inoculation with *A. radicis* resulted in microbe-induced resistance against aphids, but the strengths differed with timing of inoculation in our experiment. When bacterial inoculation was treated as one four-level factor, only the post-heatwave levels – which were closer to the moment of aphid infestation – showed significant microbe-induced resistance against aphids. However, when pre- and post-heatwave were treated as two independent inoculations in a separate model, we observed that both inoculations caused significant microbe-induced resistance against aphids, but that the post-heatwave inoculation was much stronger (Table S1). This strongly suggests that microbe-induced resistance in plants might wane over time after inoculation (Coy et al., 2019; Gadhave et al., 2016b). This has important practical implications, as it indicates that bacterial inoculation should be timed close to aphid infestation to optimize plant responses to herbivores. Our results indicate that beneficial microbial inoculations may be needed to be applied during field seasons, with restorative or potentially boosting beneficial effects, offering a promising application of beneficial microbes as environmentally friendly alternatives to suppress pests in barley crops. It should be clear that *A. radicis* inoculation alone will not change entire agricultural schemes, but it could be one of many necessary steps towards more sustainable agricultural practices. Future studies applying beneficial microbes in natural settings and using a broader range of cultivars will help us understand how beneficials operate under variable abiotic and biotic conditions, which will be an important next step toward the ecological intensification of agriculture (Zytynska, 2021).

Highlighting the consistency of microbe-induced resistance against aphids, our results show that *A. radicis* effects on aphids were not significantly altered by heatwave conditions or water availability. This is important as it indicates that microbial inoculation can achieve stable microbe-induced resistance across a wide range of climatic contexts. Although we did not measure defence responses in this study, it was shown in recent studies that microbial inoculation with *A. radicis* activates plant defences, particularly upregulating pathogenesis-related genes and flavonoid biosynthesis genes involved in phloem-feeding herbivore defences, allowing inoculated plants to have a faster and more robust defence response upon insect attack compared to uninoculated plants (Han et al., 2016; Sanchez-Mahecha et al., 2022). Similar pathways have been shown to invoke resistance against phloem-feeders in rice inoculated with *Bacillus velezensis* (Rashid et al., 2018).

Various plant defenses have been shown to interact with abiotic conditions. For instance, under high temperatures, plants can activate biochemical responses, such as producing reactive molecules like reactive oxygen species to cope with thermal stress (Lamaoui et al., 2018; Yoshida et al., 2014) that are also involved in plant defenses against herbivores (Hillwig et al., 2016; Hirayama & Shinozaki, 2010). Contrary to our expectations, heatwaves did not leave long-lasting effects that affected aphid colony growth. However, on the one hand, the insects never had direct exposure to the heatwave, a climatic event that has been shown to negatively impact insect performance directly (Beetge et al., 2019; Ma et al., 2004; Nguyen et al., 2009). On the other hand, it could be that the abiotic conditions were not extreme enough to result in lasting physiological changes in plants. This might be because even the most stressful abiotic conditions (i.e., minimally-watered heatwave exposed plants) were still watered, albeit at a low level. Although little is known about long-lasting physiological effects (i.e., legacy effects) of extreme climatic events on plant resistance (Harvey et al. 2020), our study suggests that temperature extremes leave no measurable, lasting effects on insect resistance in our model system.

*Acidovorax radicis* had a consistent suppressive effect on aphids across all abiotic conditions; this is contrary to our hypothesis that heatwaves and low water availability conditions would disrupt the microbial-plant interaction and that a reinoculation could restore this interaction and its subsequent effects on microbe-induced resistance. One plausible explanation might be that *A. radicis* forms biofilms that increase the odds that the bacteria colonize the plant and make a beneficial association (Ramey et al., 2004). Bacterial biofilm formation has been associated with increased water retention in other bacterial species and helps increase the plant-bacterial beneficial interactions (Timmusk & Nevo, 2011; Valliere et al., 2020), and therefore may play an important role in providing stability under varying abiotic environments (i.e., providing protection against extremes in water availability during heatwave periods). The consistency of microbe-induced resistance also falls in line with our general observation that aphid colony growth overall was similar across climatic treatments. This reinforces our conclusion that temperature extremes did not leave lasting physiological changes in our model plants.

### *Acidovorax radicis* minimally affects plant growth but interacts with climate factors

Bacterial inoculation did not show any main effects on plant growth or chlorophyll levels in the current study. However, we did observe an interactive effect between bacterial inoculation and heatwave treatment on chlorophyll content depended on the heatwave exposure. When plants were under ambient conditions, post-heatwave inoculation increased the chlorophyll content compared to no inoculated plants, but this positive effect for the plant disappeared under heatwave conditions. This suggests that bacterial inoculation can benefit plants by increasing photosynthetic capacity under normal conditions (Liu et al., 2019; Vishnupradeep et al., 2022) but that the effects likely wane off after a period of time post-inoculation (Sanchez-Mahecha et al., 2022). A later inoculation leads to higher chlorophyll, although this is less pronounced in the double inoculation. We speculate that the interaction between bacteria and plants, despite invoking microbe-induced resistance and plant responses in the short term, also seems to limit plants in terms of plasticity in the longer term (Goh et al., 2013). It could be that the plants perceive the bacteria as pathogens (invoking SAR-type defence responses), and invest in energy (i.e., elevated chlorophyll) to fight them off. The *Acidovorax* genus has pathogenic members known to cause disease in many other plant systems, including maize, rice, and cucumber (Siani et al., 2021), and despite not showing pathogenic symptoms in our model system, may still be recognized as such via PAMPs or other antigenic patterns. This would also explain why *A. radicis* is typically eradicated from the roots within few days (Sanchez-Mahecha et al., 2022). Heatwaves often restrict plant responses to bacteria, potentially because plants are recovering from heat stress or preserving energy for other processes (Zandalinas et al., 2017); other research has shown comparable impacts of bacteria, combined with biotic stressors leading to reduced chlorophyll content and increased stress tolerance (Rashid et al., 2017). Future work into plasticity of pathways that govern microbe-plant-insect interactions and how they function under biotic and abiotic stress conditions is urgently needed.

## Conclusion

Plant inoculation with *A. radicis* rhizobacteria invokes consistent microbe-induced resistance against aphids across a range of climatic conditions. This is important as it indicates that these microbial agents could be applied in variable conditions with stable effects. A more critical aspect appears to be timing of inoculation relative to pest arrival. To avoid waning of microbe-induced resistance, it will be essential to establish optimal time points and methods for inoculation relative to pest infestations, which requires incorporating knowledge of temporal pest dynamics in field microbial applications. This study contributes to a better understanding of how beneficial microbe-plant interactions are affected by climatic extremes. Our study illustrates that beneficial bacteria can be an important biological resource that could be a layer of protection in sustainable agricultural practices, even under the more frequent climate abnormalities that are predicted in future climate change trajectories. Effects of climatic extremes should be studied in combination with other climate change factors as climatic extremes rarely occur independently of other factors (Rillig et al., 2019), and more realistic scenarios in studies of global change are urgently needed. Follow-up experiments should aim to understand how microbe-plant-insect interactions operate under natural conditions where multiple global change drivers might influence these interactions at the same time.

## Supporting information

Suppemental 1

## Acknowledgements

We thank Zoë A. Pfeiffer and Nafiseh Mahdavi for the help during the experiment performance and Monika Plaga for aphid colony maintenance. We thank Roman Meier for assistance in the heatwave program simulation and support at TUM*mesa*. This work was funded by a grant of the Deutsche Forschungsgemeinschaft (DFG), grant numbers RO 2340/4-1 and WE 3081/36-1.

## Author contributions

OS and RH designed the experiment, with input from all co-authors. OS, RH, SK and SS performed the study and collected the data. SK prepared the bacterial inocula. OS analysed and interpreted the data with the support of RH and SZ. OS drafted the manuscript with regular input from RH and SZ. All authors have read and contributed and approved the final manuscript.

## Data availability statement

Data will be stored in an online data repository upon publication.

