## Supplementary material for "Microbe-induced plant resistance against insect pests depends on timing of inoculation, but is consistent across climatic conditions": Suppemental 1

### Overview Supporting Information

#### Figures

Figure S1. Rhizobacterial *Acidovorax radicis* effect on aphid numbers across abiotic treatments and inoculation timing.

#### Tables

Table S1. Summary of standard linear models of the plant and aphid response variables.

Table S2. Summary of model selection based on AIC values.

### Supporting Information Figure S1

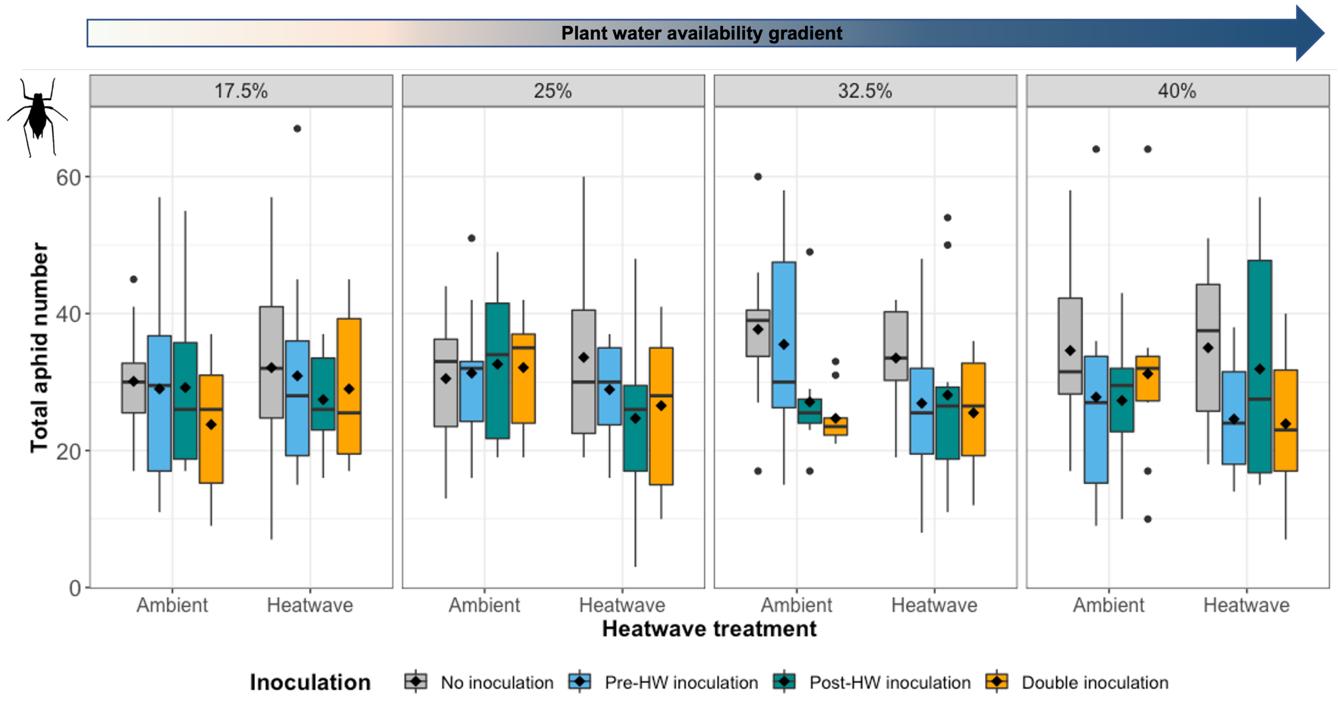


**Figure S1.** Effect of the heatwave, plant water availability and timing of rhizobacterial inoculation (*Acidovorax radicis)* on total aphid numbers (*Sitobion avenae*).

**Table S1. Summary of standard linear models for the plant and aphid response variables**. Abbreviations are as follows: Heatwave (H), Water availability (W), Pre-Heatwave bacterial inoculation (I1) and Post- Heatwave bacterial inoculation (I2). *Note:* Data were transformed as necessary to complete normality assumptions ([root biomass^0.04]; [log(shoot-to-root ratio+1]; [chlorophyll content ^2]). "Bold values" show significant p values (<0.05), from minimal adequate models. Dashes indicate terms removed after model simplification.

| | Shoot biomass | | | Root biomass | | | Shoot/root biomass ratio | | | Chlorophyll content | | | Aphid number | | |
|---|---|---|---|---|---|---|---|---|---|---|---|---|---|---|---|
| | d.f. | *F* | *p* | d.f. | *F* | *p* | d.f. | *F* | *p* | d.f. | *F* | *p* | d.f. | *F* | *p* |
| Heatwave (H) | 1 | **7.94** | **0.005** | 1 | **18.93** | **<0.001** | 1 | 0.65 | 0.422 | 1 | 1.30 | 0.255 | 1 | 1.94 | 0.165 |
| Water availability (W) | 1 | **14.07** | **<0.001** | 1 | **12.25** | **0.001** | 1 | **12.89** | **<0.001** | 1 | **13.35** | **<0.001** | 1 | 0.10 | 0.755 |
| Pre- Heatwave inoculation (I1) | 1 | 2.20 | 0.139 | 1 | 1.21 | 0.272 | 1 | 1.30 | 0.254 | 1 | 0.13 | 0.719 | 1 | **4.84** | **0.029** |
| Post- Heatwave inoculation (I2) | 1 | 1.22 | 0.270 | 1 | 0.99 | 0.320 | 1 | 0.81 | 0.370 | 1 | **7.18** | **0.008** | 1 | **8.21** | **0.004** |
| H x W | 1 | - | - | 1 | - | - | 1 | 0.15 | 0.701 | 1 | **3.99** | **0.047** | 1 | - | - |
| H x I1 | 1 | - | - | 1 | - | - | 1 | **3.98** | **0.047** | 1 | 0.67 | 0.412 | 1 | - | - |
| W x I1 | 1 | - | - | 1 | - | - | 1 | 0.48 | 0.487 | 1 | - | - | 1 | - | - |
| H x I2 | 1 | - | - | 1 | - | - | 1 | 0.26 | 0.608 | 1 | **7.01** | **0.009** | 1 | - | - |
| W x I2 | 1 | - | - | 1 | - | - | 1 | 1.38 | 0.241 | 1 | - | - | 1 | - | - |
| I1 x I2 | 1 | - | - | 1 | - | - | 1 | 0.18 | 0.674 | 1 | - | - | 1 | - | - |
| H x W x I1 | 1 | - | - | 1 | - | - | 1 | 3.72 | 0.055 | 1 | - | - | 1 | | |
| H x W x I2 | 1 | - | - | 1 | - | - | 1 | 0.31 | 0.577 | 1 | - | - | 1 | - | - |
| H x I1 x I2 | 1 | - | - | 1 | - | - | 1 | 3.62 | 0.058 | 1 | - | - | 1 | - | - |
| W x I1 x I2 | 1 | - | - | 1 | - | - | 1 | 0.00 | 0.992 | 1 | - | - | 1 | - | - |
| H x W x I1 x I2 | 1 | - | - | 1 | - | - | 1 | **4.29** | **0.039** | 1 | - | - | 1 | - | - |
| Residuals | 306 | | | 306 | | | 295 | | | 307 | | | 308 | | |

**Table S2. Model selection based on selection of AIC values, after model simplification step.**

| AIC | Shoot biomass | Root biomass | Shoot/root biomass ratio | Chlorophyll content | Aphid number |
|---|---|---|---|---|---|
| Before simplification (No selected model) | AIC=-2044.1 | AIC=-2487.01 | AIC=-840.99 | AIC=1046.02 | AIC=1540 |
| **After simplification (Selected model)** | **AIC=-2064.1** | **AIC=-2495.49** | **AIC=-845.85** | **AIC=1033.09** | **AIC=1526.04** |
